# Hairpin-locker mediated CRISPR/Cas tandem system for ultrasensitive detection of DNA without pre-amplification

**DOI:** 10.1101/2024.03.05.583466

**Authors:** Fei Deng, Rui Sang, Yi Li, Biyao Yang, Xiwen Zhai, Ruier Xue, Chengchen Zhang, Wei Deng, Ewa M. Goldys

## Abstract

Achieving ultra-sensitive detection of DNA is of paramount importance in the field of molecular analytics. Conventional amplification technologies such as polymerase chain reaction (PCR) currently play a leading role in ultrasensitive DNA detection. However, amplicon contamination common in these techniques may lead to false positives. To date, CRISPR-associated nucleases (type V & VI) with their programmable cleavage have been utilised for sensitive detection of unamplified nucleic acids in complex real samples. Nevertheless, without additional amplification strategies, the pM range sensitivity of such CRISPR/Cas sensors is not sufficient for clinical applications. Here, we established a hairpin-locker (H-locker) mediated Cas12-Cas13 tandem biosensing system (Cas12-13 tandem-sensor) for ultrasensitive detection of DNA targets. Without the need for any additional amplification reaction or device, this system is capable of detecting DNA at a notable 1 aM level (<1 copy/uL) sensitivity. In addition, the system was able to distinguish cancer mutations in colorectal cancer (CRC) mice. This is a significant advance for CRISPR/Cas biosensing technology offering simple, highly sensitive, and user-friendly diagnostics for next-generation nucleic acid detection.

## Introduction

Nucleic acid detection is widely utilised in a range of fields, such as medical diagnostics, food safety monitoring, and environmental assessment^1, 2^. The programmability of CRISPR/Cas Ribonucleoprotein (RNP) via Watson-Crick base pairing of guide RNA (gRNA) with molecular targets offers an emerging approach for nucleic acid testing with ultrahigh selectivity^3^. These RNPs require only gRNA replacement to target specific nucleic acid sequences^3^. The programmable *trans*-cleavage of Type V and VI Cas proteins makes it possible to develop sequence-specific CRISPR/Cas sensors,^3-5^ which upon target detection cleave single strand DNA/RNA reporters in classic sensor designs such as DETECTR^3^, SHERLOCK^4^, and HOLMES^6^. They are capable of detecting target nucleic acids with high specificity down to a level of single nucleotide differences. In order to reach the sensitivity level of the gold standard method polymerase chain reaction (PCR) or loop-mediated isothermal amplification (LAMP), the CRISPR/Cas sensor systems need to be integrated with nucleic acid amplification which carries risk of amplicon contamination leading to false-positives^7^.

Amplification-free detection of nucleic acids by CRISPR/Cas sensors requires a combination with other techniques and approaches such as SERS^8^, metal-enhanced fluorescence^9^, nanoenzymes^10^, and field-effect transistors^11^ which require specialised instrumentation or consumables. The Cas tandem biosensing systems such as Cas13 with Csm6 tandem system ^12^, Cas13 with Cas14 tandem system ^13^, and Cas13 with Cas12 tandem system^14^ offer a strategy for disrupting the one-to-one correspondence between target molecules and activated RNPs to realize one-to-more correspondence. However, all the current tandem systems are designed for the detection of RNA targets. Bearing this in mind, we propose here a novel tandem biosensor design to realise ultrasensitive and rapid detection of DNA.

Our CRISPR/Cas tandem biosensing system employs a specially designed nucleic acid structure in form of an asymmetric hairpin (H locker, Figure 1), which acts as a bridge between two Cas RNPs. In our approach, the activation of the first Cas RNP (Cas12) by a DNA target, allows the H-locker to be cleaved which releases a new activation trigger for the second Cas RNP (Cas13). This sensor system has been able to detect DNA at 1 aM level with the detection range spanning 11 orders of magnitude. Moreover, it successfully identified the PIK3CA-H1047R mutation in plasma samples from normal and colorectal cancer (CRC) mice.

**Figure 1.**
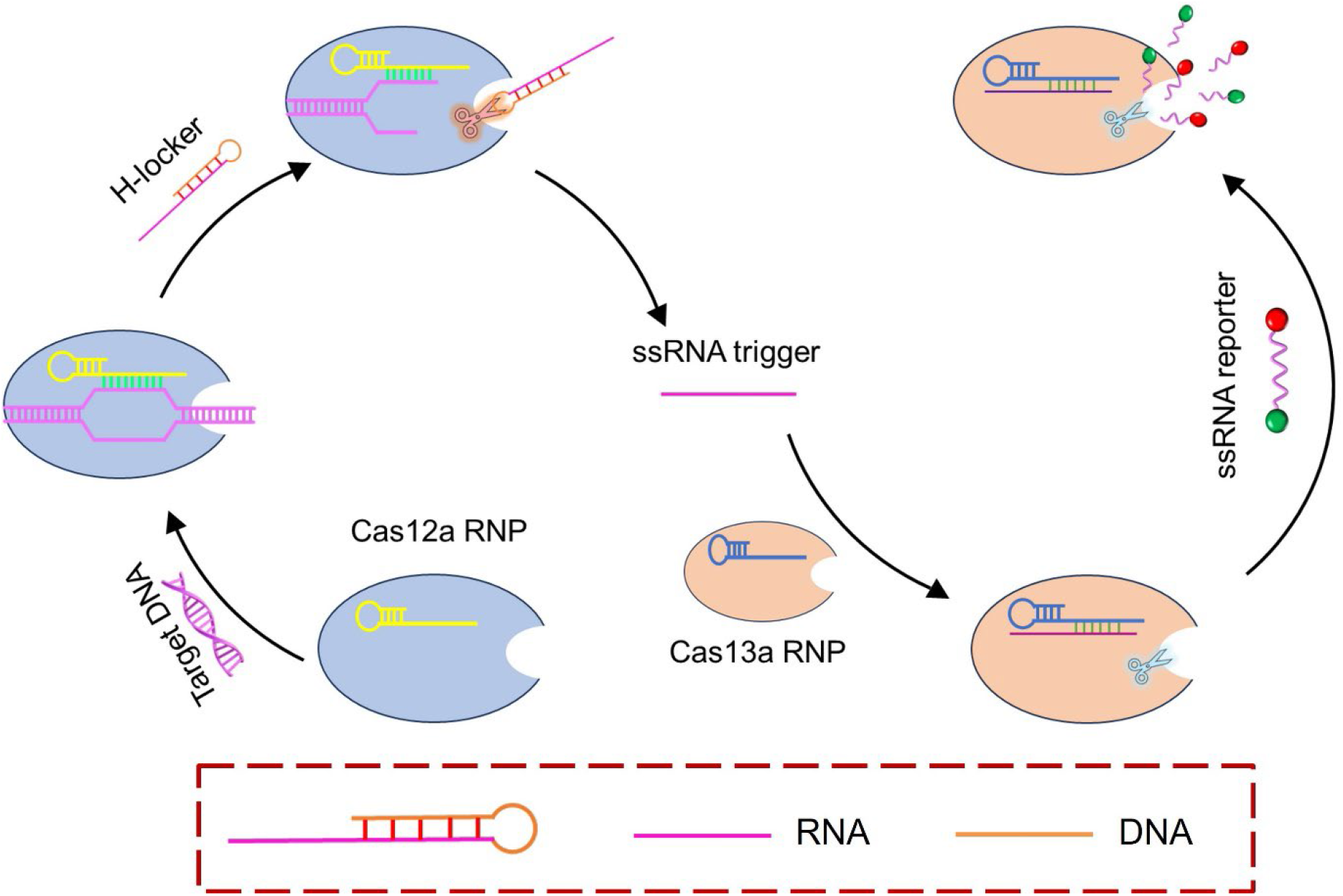
Schematic of H-locker mediated CRISPR/Cas tandem biosensing system. Upon binding to target DNA, activated Cas12a RNP *trans*-cleaves the H-locker to release the ssRNA activation trigger for Cas13a RNP, leading to a cascade signal amplification. The H-locker contains two parts, an ssRNA trigger (pink), and an ssDNA locker (orange).

## Materials and Methods

### 1. Materials and reagents

EnGen® Lba Cas12a (Cpf1) protein (New England Biolab), ENB2.1 buffer (New England Biolab), DNase/RNase free water (ThermoFisher), LwCas13a (Magigen), rCutSmart Buffer (New England Biolab), Cas13a reaction buffer (Magigen), dithiothreitol (DTT, Sigma), agarose (ThermoFisher), SYBR Gold DNA dye (ThermoFisher), 10 bp DNA ladder (ThermoFisher), 6X DNA loading dye (ThermoFisher), and phosphate buffered saline (PBS) (Sigma, 10 mM, pH=7.4).

All DNA and RNA oligos are synthesized and modified by Sangon Bio-Tech Ltd.

**Table 1.**
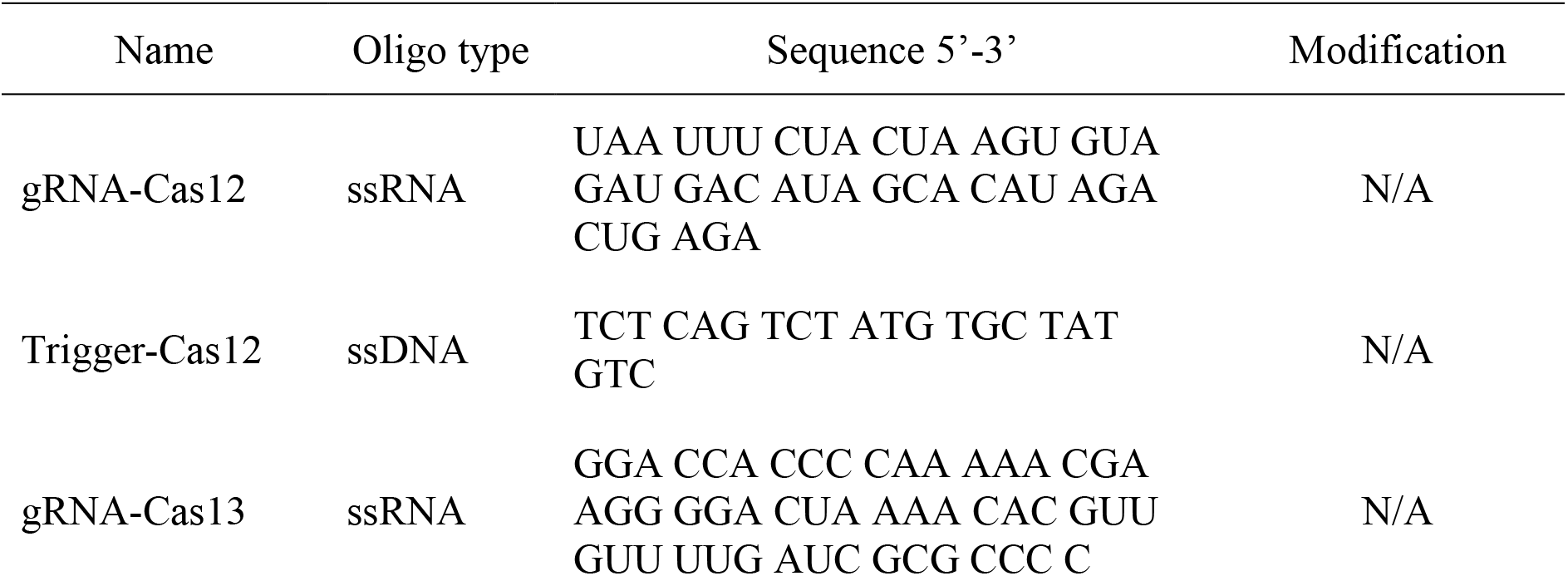
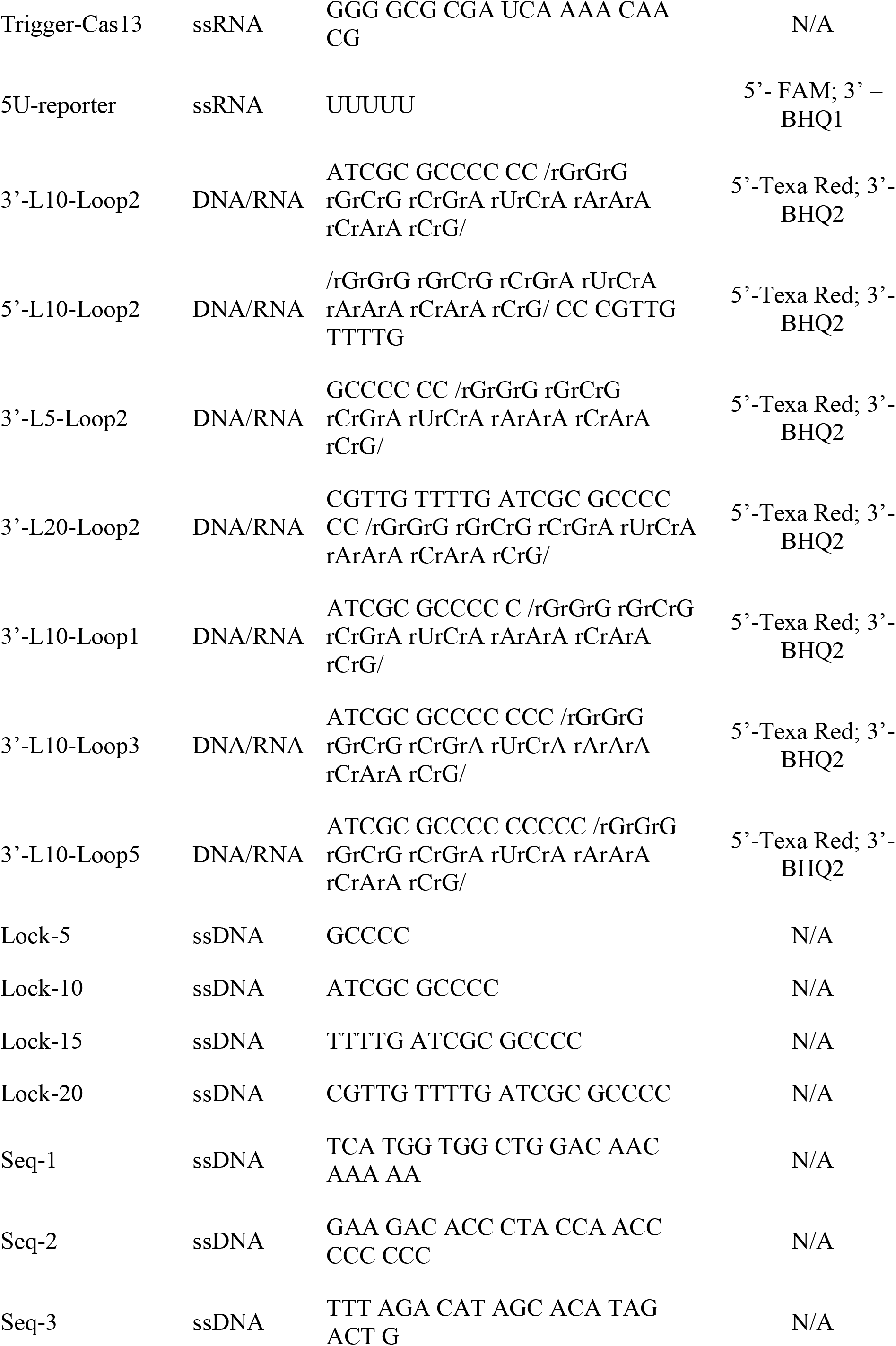
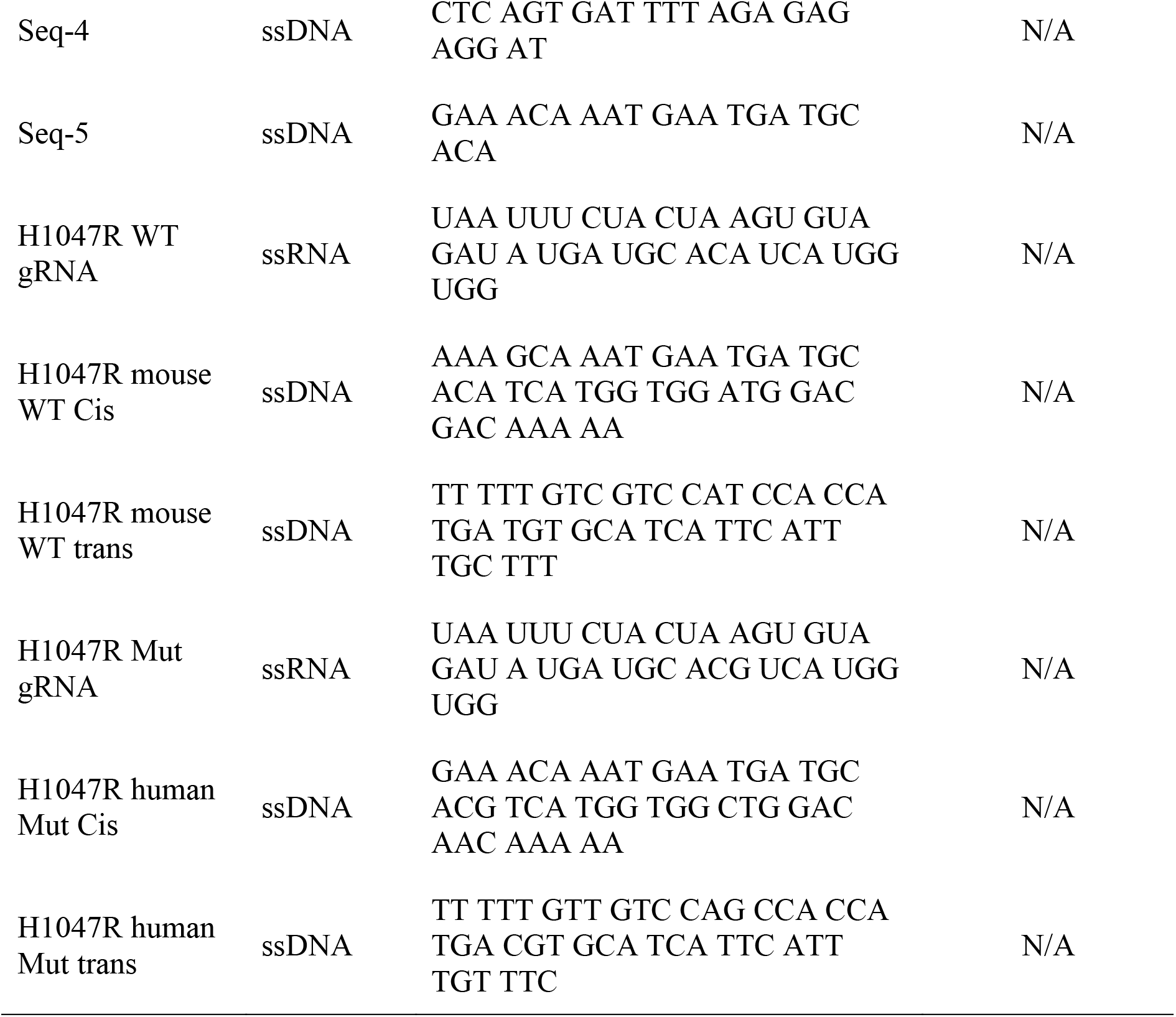
DNA and RNA oligos used in this study.

### 2. Investigation of the activation of Cas13a RNP by the H-locker

The standard CRISPR/Cas13a reaction mixture was prepared by combining 40nM of Cas13a protein, 20nM of gRNA-Cas13, and 120nM of the 5U-reporter in 1mL of the rCutSmart buffer. Subsequently, 2μL of 1μM of H-lockers was added into 100μL of the standard reaction mixture and incubated at 37°C for two hours. The fluorescence signal was tested using an ID5 plate reader (λex: 480 nm, and λem: 520 nm).

### 3. Investigation of the *trans*-cleavage ability of Cas12a on H-locker.

The Cas12a-cleaved H-locker was first prepared. In brief, 200 nM of Cas12a protein, 200 nM of gRNA-Cas12, 1 μM of the H-locker, and 200 nM ssDNA trigger were added into 100 μL of the 1X rCutSmart buffer. The reaction mixture was incubated at 37°C for two hours. Subsequently, gel electrophoresis assay was carried out. In brief, 5% agarose gel in 1X TBE buffer was prepared with SYBR Green RNA dye. 10 μL of pristine H-locker and Cas12a cleaved H-locker were premixed with 2 μL of 6X RNA gel loading dye and then loaded into the gel for electrophoresis, which was carried out for 40 min at a constant voltage of 100V. 5 μL of 10 bp DNA ladder was used for molecular weight reference. Gel images were visualized by using Gel Doc + XR image system (Bio-Rad Laboratories Inc., USA).

Subsequently, a fluorescent assay was carried out. A standard CRISPR/Cas12a reaction solution was prepared where 1 μL of 100 μM of Cas12a protein, 5 μL of 20 μM of gRNA-Cas12, and 6 μL of 100 μM of H-locker were added into 3.6 mL of 1X rCutSmart buffer. The prepared solution was stored at 4 °C before use. To further enhance the *trans*-cleavage rate, 10 mM of DTT was added in the reaction solution. Subsequently, 5 μL of 1 μM of trigger-Cas12 ssDNA was added to trigger the CRISPR/Cas12a reaction and incubated at room temperature for two hours. The fluorescence signal was tested using a SpectraMax iD5 multi-Mode Microplate Reader (λex: 570 nm; λem: 615 nm).

### 4. Investigation of the Cas13 activation by cleaved H-locker.

The standard CRISPR/Cas13a reaction mixture was prepared by combining 40nM of Cas13a protein, 20nM of gRNA-Cas13, and 120nM of 5U-reporter in 1mL 1X rCutSmart buffer. Subsequently, 2μL of 1μM of cleaved H-locker (Method 3) or DNA/RNA hybrid trigger was added into 100μL of the standard reaction mixture and incubated at 37°C for two hours. The fluorescence signal was tested using a SpectraMax iD5 multi-Mode Microplate Reader (λex: 480 nm, and λem: 520 nm).

### 5. Investigation of the biosensing performance of H-locker mediated CRISPR/Cas tandem biosensing system

The standard reaction solution was prepared as follows. In brief, 1 μL of 100 μM of Cas12a protein, 5 μL of 20 μM of gRNA-Cas12, 50 μL of 2 μM of Cas13a protein, 5 μL of 20 μM of gRNA-Cas13, 15 μL of 20 μM of 5U-reporter, and 36 μL of 1 M of DTT were added into 3.6 mL of rCutSmart buffer. The prepared solution was stored in 4 °C before use.

To assess sensitivity, different concentrations of trigger ssDNA and 3 μL of 1 μM of H-locker were added to 100 μL prepared reaction solution to trigger the tandem system and incubated at 37°C for two hours. The fluorescence signal was tested using SpectraMax iD5 multi-Mode Microplate Reader (λex: 480 nm, and λem: 520 nm).

To assess specificity, 10 nM of trigger ssDNA or interference DNA sequences and 3 μL of 1 μM of H-locker were added to 100 μL of the prepared reaction solution to trigger the tandem system and incubated at 37°C for two hours. The fluorescence signal was tested using a SpectraMax iD5 multi-Mode Microplate Reader (λex: 480 nm, and λem: 520 nm).

To assess stability, 120nM of fluorescent labelled H-locker was incubated in 0, 2%, 5%, 10% mouse plasma/rCutbuffer solution for two hours. In addition, to eliminate the impact of plasma nuclease, 10% mouse plasma/rCutbuffer solution was first heat treated at 95°C for 10 min and 30 min, subsequently, 120nM of fluorescent labelled H-locker was added into the solution and incubated for two hours. The fluorescence signal was tested using a SpectraMax iD5 multi-Mode Microplate Reader (λex: 480 nm, and λem: 520 nm).

### 6. The application of tandem sensor for the detection of ctDNA from mouse plasma samples

For the detection of ctDNA from mouse plasma samples, a standard reaction solution was prepared. In brief, 1 μL of 100 μM of Cas12a protein, 5 μL of 20 μM of H1047R Mut gRNA or H1047R WT gRNA, 50 μL of 2 μM of Cas13a protein, 5 μL of 20 μM of gRNA-Cas13, 15 μL of 20 μM of 5U-reporter, and 36 μL of 1 M of DTT were added into 3.6 mL of 1X rCutSmart buffer. The prepared solution was stored at 4 °C before use. The mouse plasma samples were first diluted in rCutSmart buffer at 1:1 ratio, then subjected to heat treatment at 95°C for 30 min before use. Afterwards, 10 μL of heat-treated mouse plasma samples were added into 100 μL of the tandem reaction solution to trigger the reaction and incubated at 37°C for two hours. The fluorescence signal was tested using a SpectraMax iD5 multi-Mode Microplate Reader (λex: 480 nm, and λem: 520 nm).

## Results

### 1. Schematic of H-locker mediated CRISPR/Cas tandem biosensing system

The H-locker mediated CRISPR/Cas tandem biosensing system is designed as shown in Fig 1. The H-locker contains two parts, a ssRNA part as the activation trigger for Cas13a, and a ssDNA part as the locker. Target detection is initiated by a target DNA to activate the Cas12a RNP. Subsequently, the activated Cas12a RNP cleaves the loop region of H-locker to release the ssRNA trigger, leading to the activation of Cas13a RNP. The activated Cas13a RNP further cleaves the ssRNA reporters giving rise to signal amplification. In comparison with the standard CRISPR/Cas system where one target only activates one Cas RNP, in our tandem system, one target activates one Cas12a RNP, which cleaves numerous H-lockers, leading to the activation of numerous Cas13a RNPs.

### 2. Restricted activation of Cas13a RNP by the H-locker

To establish the H-locker-mediated CRISPR/Cas tandem system, the ability of the H-locker to activate Cas13a RNP was first investigated. As shown in Fig. 2A, with the increase of incubation time, the fluorescence signal of the H-locker slightly increased. In comparison with the fluorescence signal of standard linear ssRNA trigger, the H-locker’s signal value has been significantly reduced to 7.6% at 120 min. Therefore, the H-locker is an effective locker to block the activation of Cas13a RNP by ssRNA. Subsequently, we systematically optimized the H-locker design including the locker position, locker length, and loop size. We first investigated the locker position. As shown in Fig. 2B, the fluorescence signal of 5’-LOCK was comparable to standard ssRNA trigger, while the fluorescence signal of 3’-LOCK was 12.4% of the signal of standard ssRNA trigger. Thus, the optimum position for the locker was 3’-LOCK.

**Figure 2.**
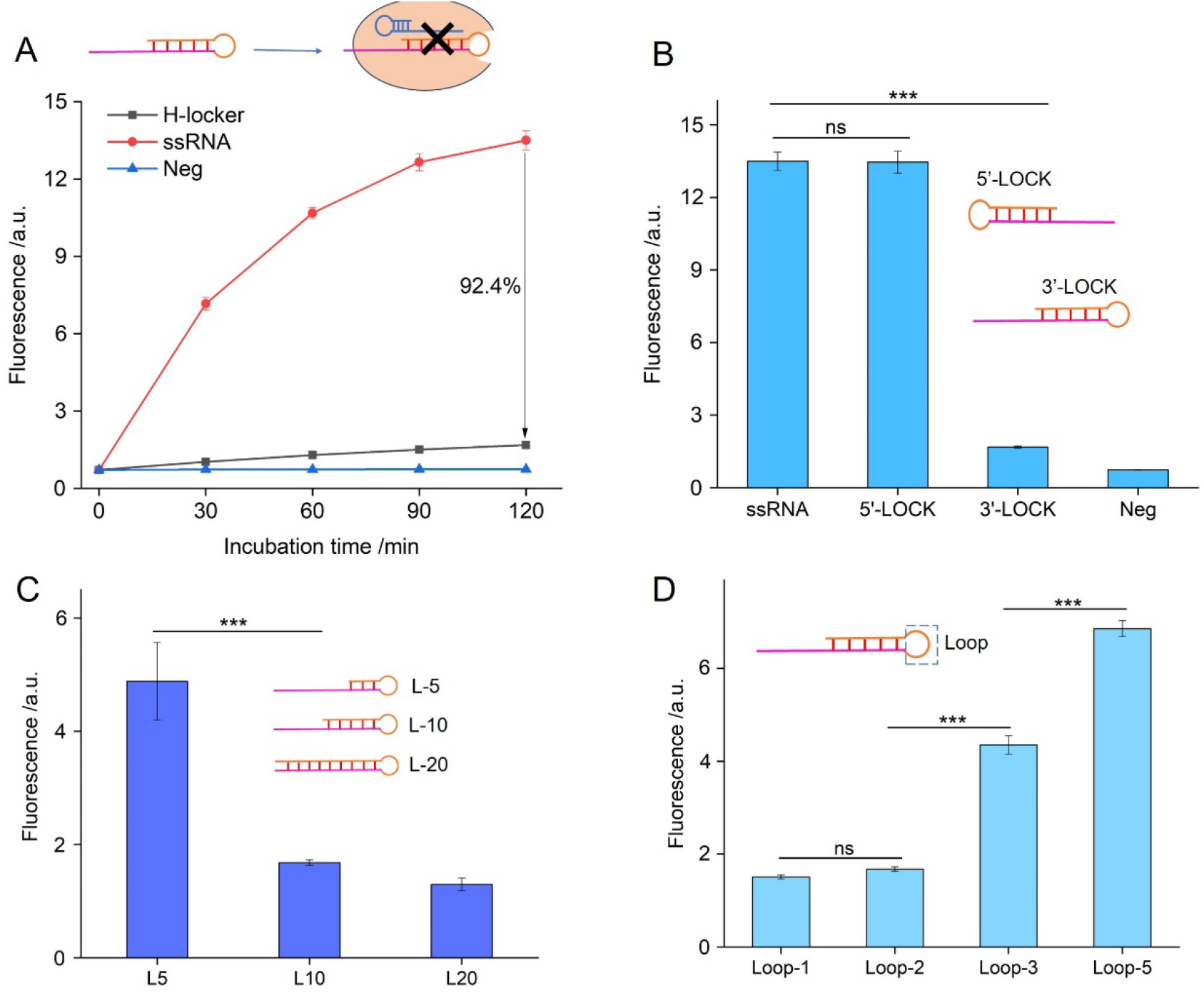
Restricted activation of Cas13a RNP by the H-locker. (A) Investigation of the activation ability of the H-locker (3’-L10-Loop2) to Cas13a RNP, and ssRNA trigger was utilized for comparison, and Neg represents no trigger (n=3); (B) Investigation of the locker position of H-locker at 5’-end or 3’-end (n=3); (C) Investigation of the locker length of H-locker, and L represents the locker (n=3); (D) Investigation of the loop size of H-locker (n=3). (* P<0.05, ** P< 0.005, *** P< 0.001)

Subsequently, the impact of locker length is investigated in Fig. 2C. We found that with the increase of locker length, lower background signal was observed, confirming that the locker effect becomes stronger at longer locker lengths. In this study, when the locker length increased to 10nt, the background fluorescence signal was reduced to 12.5% of the signal for a standard ssRNA trigger. Further increases of locker length resulted in limited background signal reduction, thus 10nt was determined to be optimal. Finally, we optimized the loop size of the H-locker (Fig. 2D). A higher background signal was observed with a larger loop, and loop size below 2 nt was demonstrated to be optimal. In addition, since larger loop is easier to be opened by activated Cas12a, therefore, optimized H-locker 3’-L10-Loop2 was used in the rest of this study.

### 3. Investigation of the *trans*-cleavage capacity of Cas12a on H-locker.

In this section, we investigated the *trans*-cleavage capacity of Cas12a on optimized H-locker (3’-L10-Loop2). Gel electrophoresis assay was first applied to evaluate the *trans*-cleavage activity of Cas12a on H-locker. As shown in Fig. 3A, the original intact H-locker shows a clear band at around 30bp, while a cleaved H-locker gives rise to two bands, one corresponding to an intact H-locker, and another due to the ssRNA region of H-locker. This data demonstrates that the H-locker can be cleaved by activated Cas12a RNP to release the ssRNA region. We further established a fluorescent assay to assess the H-locker opening process. To this aim, a fluorophore and a corresponding quencher were immobilized on both ends of the H-locker (Fig. 3B), thus, initially the fluorescence signal of H-locker was quenched. Following the activation of Cas12a RNP, the loop region of H-locker is expected to be cleaved, leading to the recovery of fluorescence signal. As shown in Fig. 3B, with the increase of incubation time, the fluorescence signal indeed continues to increase, confirming the opening of the H-locker has taken place. Finally, to enhance the Cas12a *trans*-cleavage on H-locker, a suite of previously reported chemical enhancers was applied in the reaction system^2^, and we found that DTT shows a marked enhancement effect (Fig. 3C).

**Figure 3.**
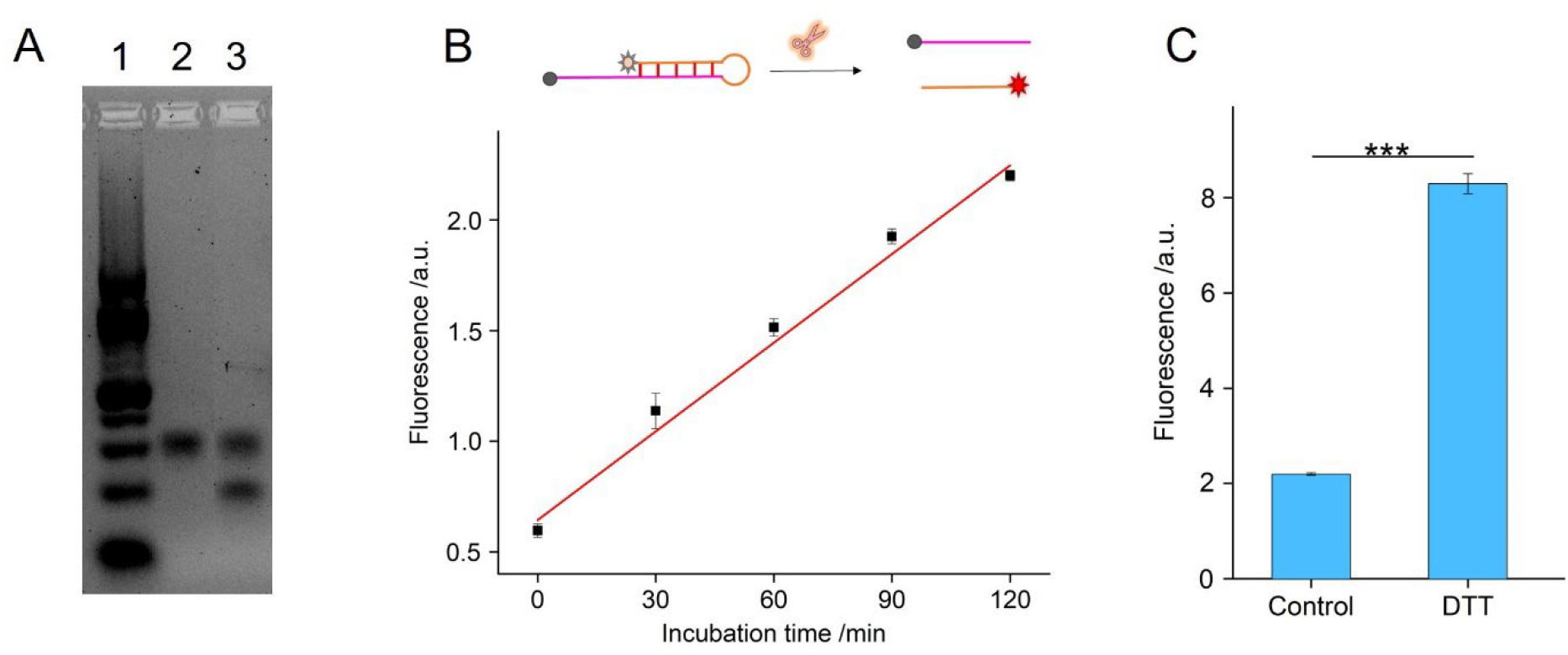
Investigation of Cas12a *trans*-cleavage on H-locker. (A) Investigation of the *trans*-cleavage activity of Cas12a on H-locker using gel electrophoresis assay. 1). 10 bp ladder; 2). H-locker; 3) Cleaved H-locker. (B) Investigation of the *trans*-cleavage activity of Cas12a on the fluorescent H-locker (n=3); (C) Enhancement effect of DTT on the *trans*-cleavage activity of Cas12a on the fluorescent H-locker (n=3). (* P<0.05, ** P< 0.005, *** P< 0.001)

### 4. Restoring the triggering ability of cleaved H-locker

In this section, we investigated whether the cleaved H-locker is able to activate Cas13a RNP. As illustrated in Fig. 4A, the fluorescence signal of a cleaved H-locker is comparable to that of the standard ssRNA trigger, demonstrating the recovery of the triggering ability of a cleaved H-locker. Additionally, to assess the impact of the ssDNA locker on the triggering ability of a cleaved H-locker, we examined the activation of Cas13a RNP by a range of DNA/RNA molecules. As depicted in Fig. 4B, a lower signal was observed with an increase in ssDNA length, and the ability of L-10 to trigger Cas13a activation was comparable to that of ssRNA. Thus, we conclude that the ssDNA locker has no significant impact on the triggering ability of a cleaved H-locker.

**Figure 4.**
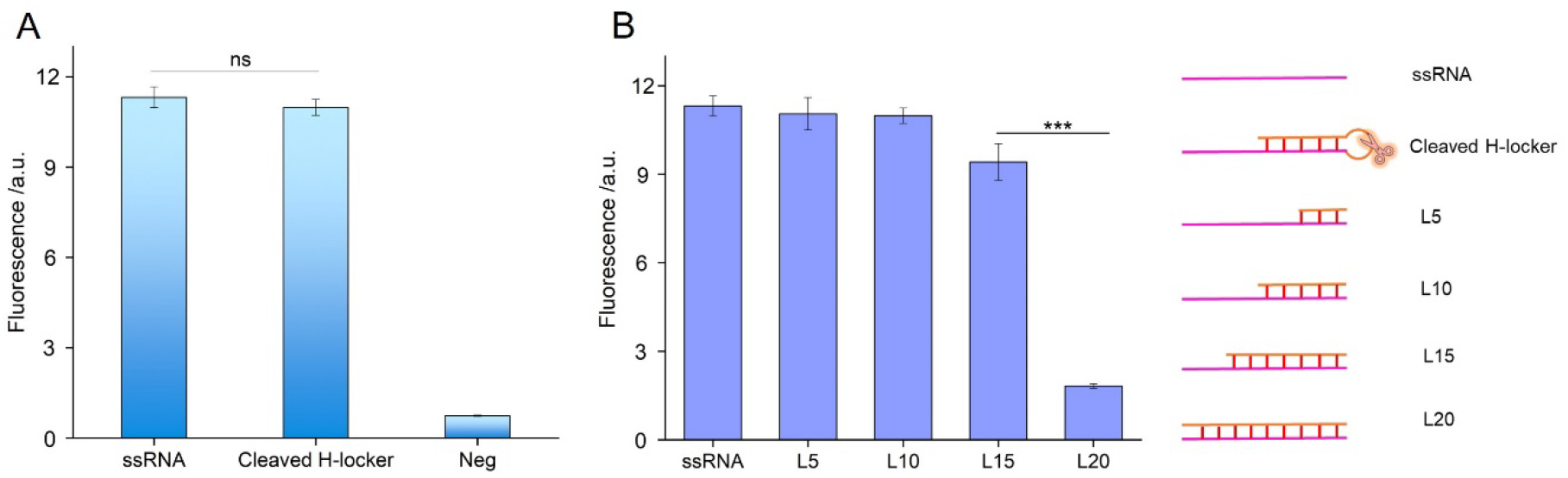
Recovery of the activation ability of H-locker to trigger Cas13a RNP. (A) Comparison of the activation ability of a cleaved H-locker and a standard ssRNA trigger to trigger Cas13a RNP (n=3); (B) Investigation of the activation ability of an RNA/DNA hybrid trigger to Cas13a RNP (n=3). (* P<0.05, ** P< 0.005, *** P< 0.001)

### 5. Investigation of the biosensing performance of H-locker-mediated CRISPR/Cas tandem biosensing system.

After evaluating the fundamental properties of H-locker, it was applied to establish the H-locker-mediated CRISPR/Cas tandem biosensing system (Fig. 1). Since both Cas12a RNP and Cas13a RNP will be expected to function in the same buffer system, the reaction buffer was first evaluated (Fig. S1), and rCutSmart buffer was found to be suitable for both RNPs. Subsequently, the sensitivity of this tandem system was evaluated (Fig. 5A). The limit of detection was determined to be 1aM with 11 logs detection range from 1aM to 10nM. Afterwards, the exceptional specificity of the tandem system was demonstrated (Fig. 5B). However, as the H-locker was not stable in mouse plasma (Fig. 5C), and a heat treatment was introduced to reduce the nuclease activity.

**Figure 5.**
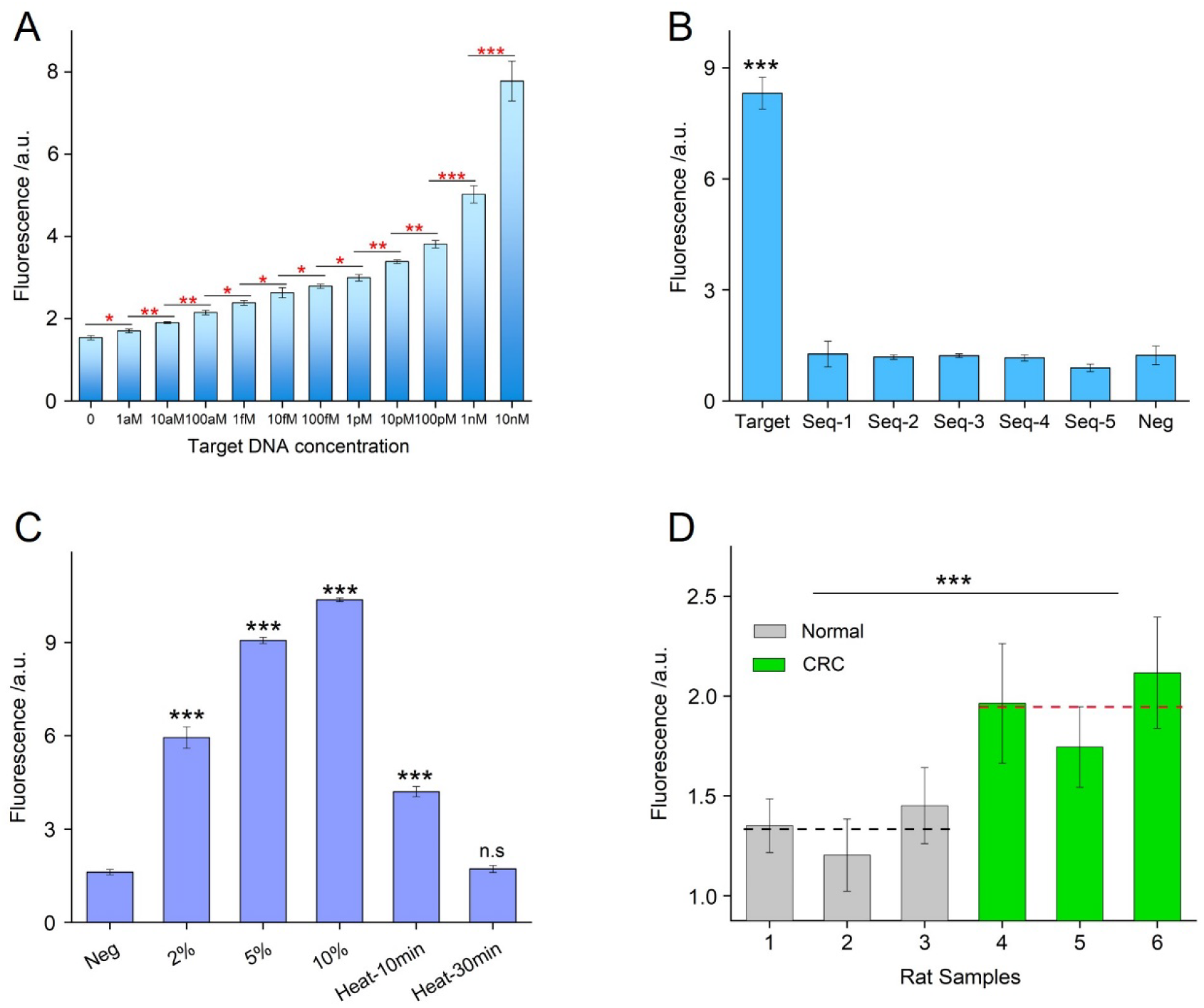
Investigation of the biosensing performance of H-locker mediated CRISPR/Cas tandem biosensing system. (A) Evaluating the sensitivity and detection range of biosensor (n=3); (B) Evaluating the specificity of biosensor (n=3); (C) Evaluating the stability of H-locker in mouse plasma (n=3); (D) Demonstrating the potential clinical applicability of biosensor for the testing of PIK3CA H1047R from CRC plasma samples (n=3). (* P<0.05, ** P< 0.005, *** P< 0.001)

Finally, the tandem system thus established was applied to detect mutant ctDNA from normal and orthotropic CRC mouse plasma samples (Fig. 5D and Fig. S2). Since the selected target sequence (19 nt) of human PIK3CA H1047R mutation and mouse PIK3CA H1047R wild type differ only by a single nucleotide (Table S1), we tested both the mutation sequence concentration by using H1047R Mut gRNA (Fig. 5D) and the wild type sequence concentration by using H1047R WT gRNA (Fig. S2). Significantly higher mutant ctDNA concentration was observed in orthotropic CRC mouse plasma samples than in healthy controls (Fig. 5D), while no significant differences were observed in the wild type concentration (Fig. S2), demonstrating the potential clinical applicability of this tandem system.

## Discussion

In this study, we developed a novel Cas12-Cas13 tandem biosensing system able to ultrasensitive detection of a single copy of DNA without amplification (Fig. 5A). Additionally, it has the ability to monitor PIK3CA-H1047R mutation from mice with human colorectal cancer xenografts (Fig. 5D). A tandem sensor is a dual amplification system, comprising Cas12a RNP, Cas13a RNP, and H-locker. H-locker as the key component has the ability to activate downstream Cas13a RNPs to produce a cascade reaction system. In addition, the H-locker is a specific DNA-RNA hybrid sequence which does not require complex synthesis procedures.

Our tandem sensor provides a new user-friendly format for ultrasensitive quantification of nucleic acids. It has the ability to overcome existing translational barriers, as it is a single pot reaction. In contrast to conventional nucleic acid amplification technologies^15^, such as PCR, RPA, and LAMP, this tandem sensor provides comparable sensitivity (1 aM) but does not require sophisticated instruments and it avoids amplicon contamination. In comparison with recently established Cas feedback circuit^16^, which requires more than four hours turnover time due to the complex system design, our tandem sensor provides an cascade system for rapid detection of 1 aM nucleic acids within 60 min.

Additionally, the signal amplification components of tandem sensor (H-locker, Cas13a RNP, and ssRNA reporter) can be directly integrated into other type V based CRISPR/Cas biosensing system (Cas12 & Cas14) as an additional signal amplifier to enhance their sensitivity, without any additional changes of the original reagents or setup.^17^ Nevertheless, The H-locker still provides a certain amount of background signal (7.6%). Thus, to reduce the background signal it is essential to separately store the H-locker from the Cas12a-Cas13a reaction mixture. In conclusion, tandem sensor provides a breakthrough approach for rapid, point-of-care, and ultrasensitive quantification of nucleic acids.

## Notes

### Competing Interest Statement

The authors have declared no competing interest.

