## Supplemental file for "Hairpin-locker mediated CRISPR/Cas tandem system for ultrasensitive detection of DNA without pre-amplification"

### Supplementary Methods

1. Establishment of orthotropic CRC mouse model

All animal experiments were approved by the UNSW Animal Care and Ethics Committee (project approval 20/95B, 21/39B, and 21/77B). 6-8-week-old NOD/SCID mice were ordered by Animal Services from the Animal Resources Centre (ARC, Perth, WA). Mice were housed in specific pathogen free conditions at 22℃ with a 12 h light/dark cycle. They were accommodated in standard ventilated cages and allowed a 7 days acclimation period upon arrival at the UNSW animal facility. Mice were provided food and water ad libitum with regular monitoring of their wellbeing.

An orthotropic CRC mouse model was developed through intra-rectal tumour cell injection method with minor modifications of the previously reported work.[1] Briefly, female NOD/SCID mice (6-8-week-old) were fasted of food for 6 h prior to cancer cell injections, followed by rapid anesthesia induction with 2-4% isoflurane and maintenance at 1-3% with 1 L/min oxygen. Blunt-tip forceps were applied to dilate the anal canal, exposing the distal anal and rectal mucosa. Subsequently, $4\times10^5$ HCT-116-Luc2 cells, suspended in 10 μL of PBS and 10 μL of Matrigel, were orthotopically inoculated into the submucosa of the distal posterior rectal, approximately 1-2 mm above the anal canal, using a 30-gauge needle (Terumo, Tokyo, Japan). Mice were closely monitored for 1 to 72 h post-injection for early detection of adverse events, with subsequent monitoring occurring at least bi-weekly.

Tumour growths were monitored every 7 days through the IVIS Spectrum CT imaging system (Perkin Elmer, Waltham, US). Typically, mice were intraperitoneally injected with 150 mg/kg of D-Luciferin. Mice were then anesthetized with isoflurane, with anaesthesia maintained throughout imaging using the IVIS spectrum imaging system for bioluminescence detection via Living Image® 4.5.2 software. When tumour reached the 100 $mm^3$ volume (equivalent to approximately $4\text{-}6\times10^{10}$ photons/s of bioluminescence signal in this study), one group of mice were treated with X-ray radiation. At 27 days post treatment, the terminal blood collection (500~750 μL per mouse) was performed by the cardiac puncture technique with 25-gauge needles. K3 EDTA tubes were used for blood samples collection, allowing the isolation of blood plasma through centrifugation (1000 × g, 10 min). The isolated mice blood plasma was stored at -80°C for further use.

### Supplementary Figures

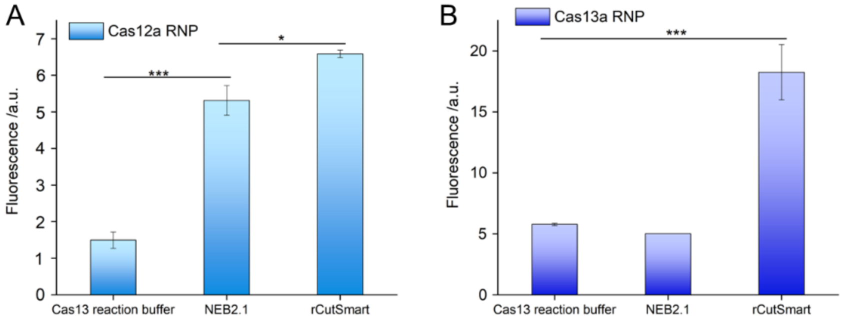


**Figure S1**. Optimization of the reaction buffer for Cas12a and Cas13a. (A) Optimization of the reaction buffer for Cas12a; (B) Optimization of the reaction buffer for Cas13a.

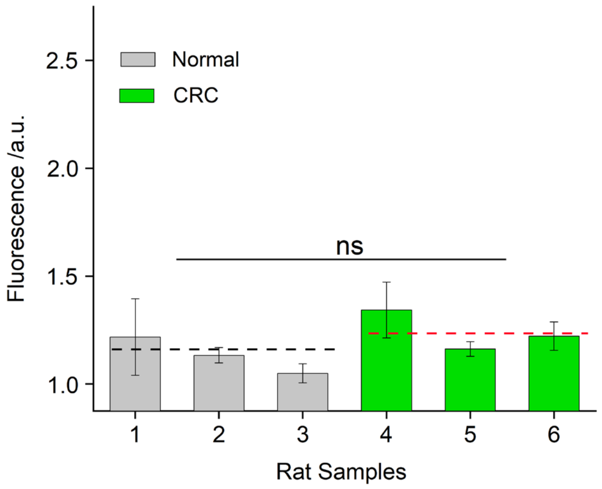


**Figure S2**. Background signal due to mouse PIK3CA H1047R wild type gene fragments in mouse plasma samples. Here, the Cas12a RNPs are targeting the wild type sequence of the mouse PIK3CA gene in the region of the H1047R mutation. The mouse PIK3CA H1047R wild type sequence differs from the human PIK3CA H1047R mutation sequence by a single nucleotide (Table S1). The fluorescence signal for the normal and CRC groups shown here are indicative of the background signal level due to the wild type PIK3CA gene fragments in the tested plasma samples. The lack of significant difference between two groups confirms that our Cas12-Cas13 tandem system can specifically detect the PIK3CA H1047R mutation in ctDNA from the mouse plasma samples, as indicated in Figure 5D. (n=3, ns = non-significant).

### Supplementary Tables

**Tables S1.** PIK3CA H1047R sequences.

| | |
|---|---|
| Human WT | 5'-GAA ACA AAT GAA TGA TGC ACA TCA TGG TGG CTG GAC AAC AAA AA-3' |
| Human Mut | 5'-GAA ACA AAT GAA TGA TGC ACG TCA TGG TGG CTG GAC AAC AAA AA-3' |
| Mouse WT | 5'-AAA GCA AAT GAA TGA TGC ACA TCA TGG TGG ATG GAC GAC AAA AA-3' |

* The red nucleic acid base is the PIK3CA H1047R mutation point. The blue section is the target sequence of Cas12a RNP, in which the human PIK3CA H1047R WT is the same as the mouse PIK3CA H1047R WT.
